# Substrate promiscuity of the *Escherichia coli* xanthine oxidase

**DOI:** 10.1101/2023.12.06.570370

**Authors:** Kristina Kronborg, Yong Everett Zhang

**Author notes:** Correspondence: Yong Everett Zhang.

## Abstract

PpnN is a cytosolic nucleosidase that cleaves nucleotide monophosphates to nucleobases and ribose-5-phosphate and plays a crucial role in regulating bacterial competitive fitness and persistence. To quantify the PpnN reaction, here we developed an enzyme-coupled assay, wherein PpnN hydrolyzes IMP to hypoxanthine, which is then converted by xanthine oxidase (XO) into xanthine, uric acid and hydrogen peroxide. The detection of hydrogen peroxide via horseradish peroxidase and Amplex Red provides a measure of PpnN enzymatic activity. Surprisingly, we found that in addition to IMP, other nucleotides like GMP significantly increased the signal, suggesting that the corresponding nucleobases might also be substrates for XO. Direct tests by using guanine confirmed XO’s capacity to use it as a substrate, albeit less effectively than hypoxanthine. These findings suggest a potential guanine aminohydrolase activity of *E. coli* XO and broaden our understanding of nucleobase metabolism in bacterial systems.

## Introduction

PpnN is a cytosolic nucleosidase involved in bacterial nucleotide metabolism (1). During stringent response, the alarmones guanosine tetra- and penta-phosphastes (p)ppGpp stimulate the enzyme activity of PpnN, facilitating a rapid adaptation of bacterial physiology to stresses (2-4). PpnN also contributes to Salmonella resistance to the host complement immune system in a (p)ppGpp dependent manner (5). However, the enzyme activity of PpnN has not been studied in detail yet.

Recently, we characterized the enzymatic activity of PpnN (2) by utilizing a mass spectrometry based method. To simplify the detection method, we explored an enzyme-coupled assay modified from (6). This assay set-up involved the initial hydrolysis of IMP by PpnN to hypoxanthine, and its subsequent conversion first to xanthine, then to uric acid and hydrogen peroxide by the xanthine oxidase enzyme (also referred to as xanthine dehydrogenase; hereafter XO). Finally, the produced hydrogen peroxide is quantitatively used via the horseradish peroxidase (HRP) to oxidize Amplex Red to resorufin, a fluorescent chemical with optimal activation at 545 nm and absorption at 590 nm (**Figure 1**). XO is present both in prokaryotes and eukaryotes and is required to produce uric acid by the breakdown of hypoxanthine and xanthine (7). The *E. coli* XO complex, composed of three proteins XdhA, XdhB, XdhC, were known to only work on the substrates hypoxanthine and xanthine according to EcoCyc and literature mining. In this study, however, we accidentally observed that XO, from SIGMA (X2252-25UN), can utilize other nucleobases as substrates as well. This suggests the potential of XO for detecting a broader range of nucleobases during enzyme assays and indicates that XO may have a weak guanine aminohydrolase activity similar to *E. coli* GuaD.

**Figure 1.**
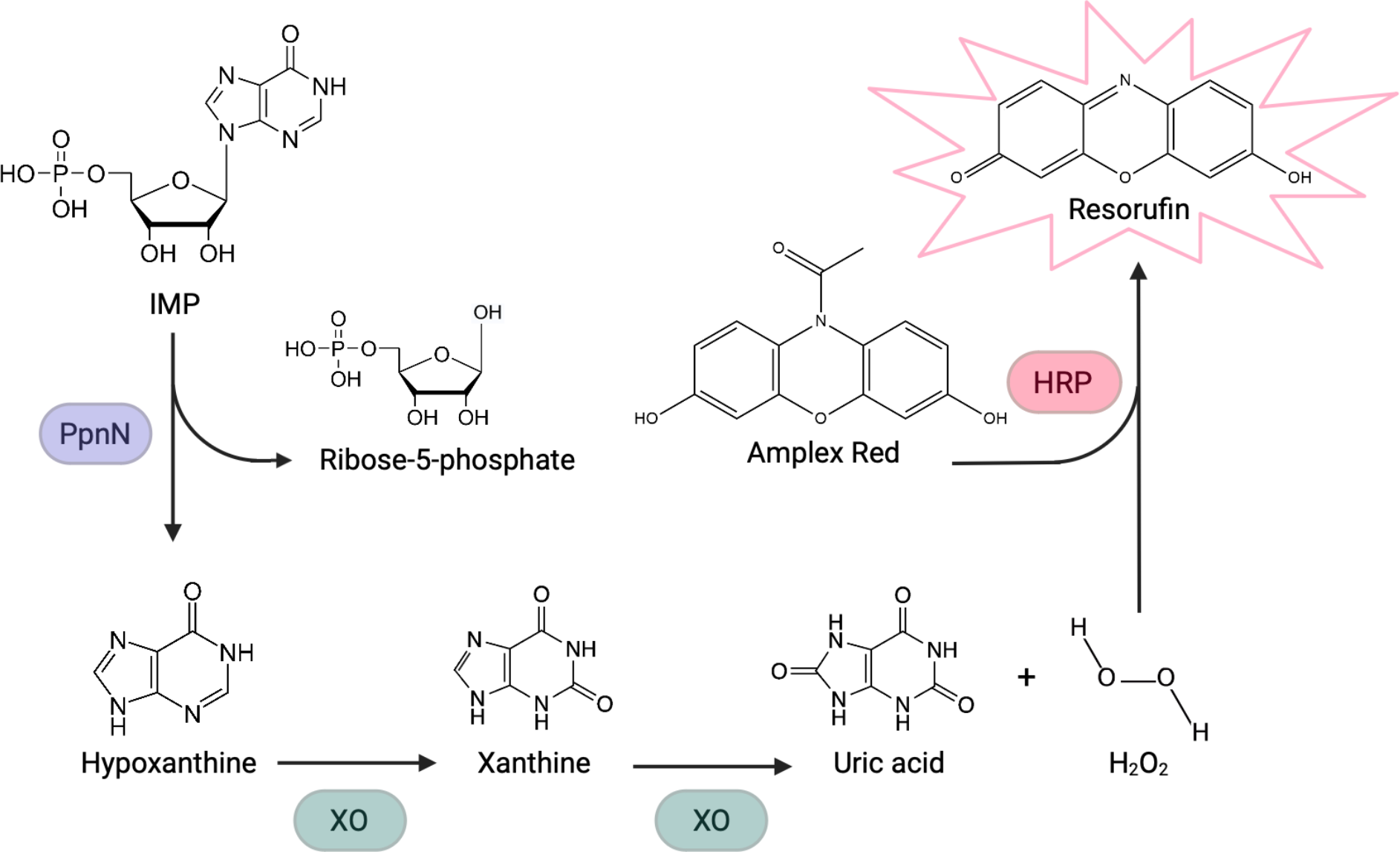
Schematic overview of the PpnN enzymatic assay set-up. First, PpnN catalyzes the hydrolysis of IMP to ribose-5-phosphate and hypoxanthine, the latter of which is used as substrate for the downstream xanthine oxidase (XO) reaction that first converts it to xanthine and then to uric acid and hydrogen peroxide. The hydrogen peroxide is finally used by horseradish peroxidase (HRP) to convert Amplex Red Reagent into a fluorescent resorufin, which is detectable spectrophotometrically at excitation- and emission coefficient of 545 and 590 nm, respectively. Figure is made using BioRender.

## Results

To determine the enzymatic activity of PpnN we initially used IMP as the substrate since hypoxanthine will be produced, which can be quantified by the XO/HRP provided by SIGMA (X2252-25UN). As we wanted to further test for any competitive effect from other nucleotides for PpnN, AMP, GMP, CMP or UMP were added to the PpnN reaction besides a fixed 5 mM of IMP. We anticipated a competitive inhibition of the PpnN’s cleavage of IMP, resulting in less hypoxanthine being produced and thus a lower final fluorescent signal. As can be seen from **Figure 2A**, however, the final signals were significantly higher in the reactions containing both IMP and another different NMP, especially for GMP, as compared to the reaction containing only IMP. Similar trends were observed for the PpnN homolog in *Salmonella typhimurium* (**Figure 2A**), despite a higher activity to IMP.

**Figure 2.**
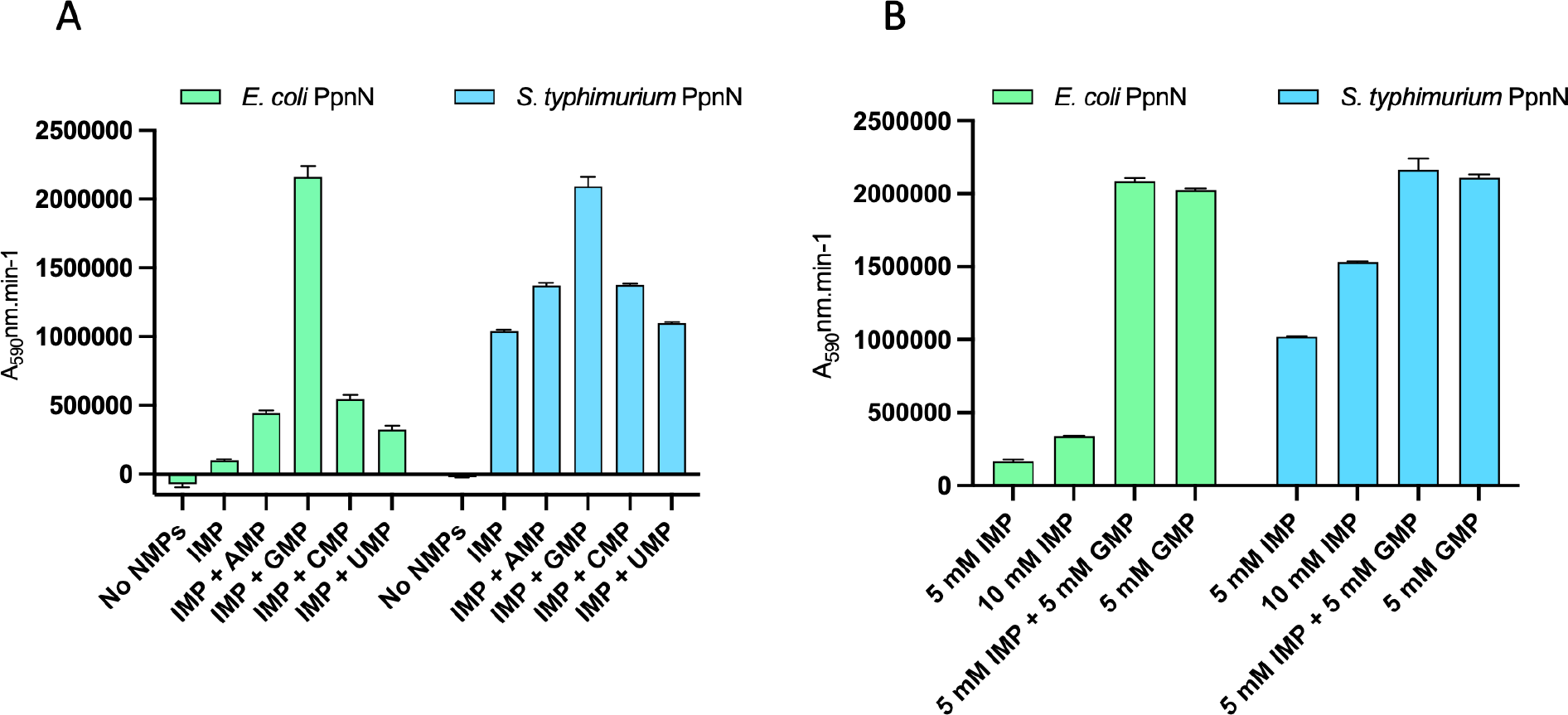
PpnN enzymatic assays in the presence of different nucleotides. Data show 30 min PpnN reactions. **A**: the presence of NMP increased the final resorufin fluorescent signal. Each NMP was used at a final concentration of 5 mM. **B**: GMP alone produced similar final fluorescent signal as the combined GMP and IMP substrates.

To better compare the reactions, we repeated the assay with reactions containing either 5 or 10 mM IMP alone, 5 mM GMP alone, or 5 mM IMP and GMP together. Extra 5 mM IMP did not boost the reaction significantly, with max. two times higher signal (**Figure 2B**, second vs the first bar). However, 5 mM extra GMP increased the signal by approx. 12 times, suggesting that either GMP stimulates the degradation of IMP or GMP itself was cleaved by PpnN to contribute to the higher signal. The former is unlikely since GMP is expected to compete for binding to PpnN given their chemical similarity. Importantly, 5 mM GMP alone also produced similar amount of signal as the combination of 5 mM GMP and IMP, demonstrating that the higher signal from the mixed GMP-IMP mainly resulted from GMP. Notably, similar trends were observed for the PpnN homolog in *Salmonella typhimurium* (**Figure 2B**). These data are consistent with the observation that GMP is a better substrate than IMP for PpnN (2). However, since XO was only known to work with hypoxanthine and xanthine, these data (**Figure 2A, 2B**) indicate that XO has extra activity towards other nucleobases, particularly guanine.

To directly confirm that XO could work on guanine, we performed the XO/HRP continuous reactions with either hypoxanthine or guanine as the substrate (**Figure 3**). This showed that XO indeed used guanine as a substrate, however, to a much lesser extent than hypoxanthine. Altogether, it is conceivable that PpnN preferentially cleaves GMP to guanine, which could be directly used by XO to generate (hypo)xanthine, H_2_O_2_ and the oxidation of amplex red to resorufin.

**Figure 3.**
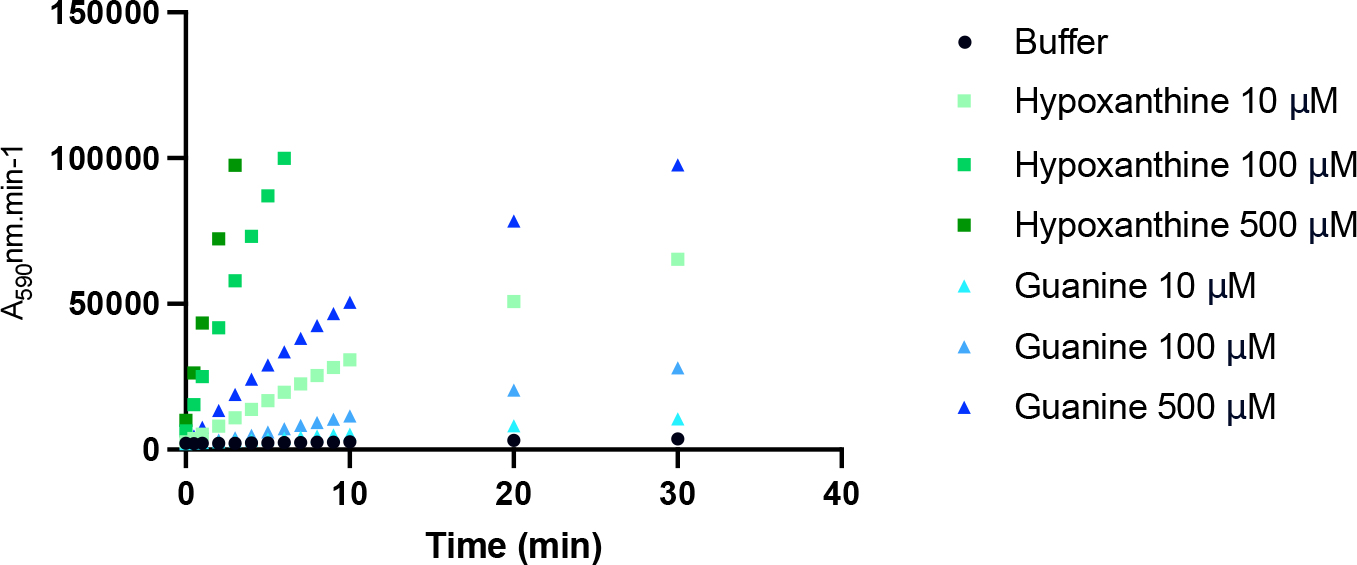
Xanthine oxidase is active on both hypoxanthine and guanine. Continuous assay testing XO activity towards 10, 100, and 500 μM hypoxanthine or guanine.

## Discussion

The results presented here indicate the activity of xanthine oxidase to be broader than initially thought. We first tested the PpnN’s ability to hydrolyze IMP in the presence of what we considered to be competing nucleotides but instead we observed a higher signal in the presence of these compounds, especially GMP. We first speculated if these nucleotides pose a stimulatory effect on the PpnN activity, hereby causing the higher signal observed in their presence. However, there was also a possibility that the xanthine oxidase used for the downstream reaction can utilize the nucleobases produced from the PpnN reaction as a substrate. Thus, we tested GMP in the presence and absence of IMP and found that GMP alone gave a high signal, suggesting that guanine can serve as substrate for xanthine oxidase. Then, we confirmed this directly by adding guanine to the xanthine oxidase/horseradish peroxidase reaction. Guanine differs from hypoxanthine by having an amino group at the C-2 position where hypoxanthine has no functional group and where a carbonyl group is added during the xanthine oxidase reaction. This could suggest a guanine aminohydrolase activity of xanthine oxidase.

As seen in **Figure 2A**, the presence of AMP, CMP or UMP also led to a higher signal compared to IMP alone, suggesting that their corresponding nucleobases can also serve as substrate for xanthine oxidase. Chemically, adenine and hypoxanthine are identical at the C-2 and C-8 that are modified by xanthine oxidase, and it is therefore not surprising that adenine might also serve as substrate for this enzyme. However, detailed further study is needed to clarify if this is the case. In particular, the XO complex needs to be reconstituted by using purified three subunits XdhA, XdhB and XdhC, besides several other cofactors, to precisely verify the mechanistic details.

## Methods

### PpnN enzymatic assay

Wild type His_6_-PpnN were purified as described in (2). All biochemical reactions were performed at room temperature. Purified PpnN proteins were first normalized to a concentration of 1500 nM in PpnN reaction buffer (25 mM HEPES pH 7.5, 130 mM NaCl, 5 mM KCl, 1.5 mM MgCl_2_ and 0.3 μM BSA) and 5 μl of this was then transferred to 40 μl PpnN reaction buffer. These 45 μl were then mixed up with 5 μl substrate to start the PpnN reaction. After selected timepoints, 10 μl of this reaction mixture was then transferred to 10 μl 1 M Na_2_CO_3_ to denature PpnN and terminate the reaction. After restoring the pH to neutral with 1 M HCl, 20 μl of this mixture was transferred to a Corning™ 96-well solid black microplate with flat bottom (10022561, Fisher Scientific) and 170 μl of a XO/HRP buffer composed of 25 mM HEPES pH 7.5, 130 mM NaCl, 5 mM KCl, 1.5 mM CaCl_2_, 1 mM MgSO_4_, 5 mM glucose, 0.3 μM BSA, 1 U/ml HRP, 60 μM Amplex™ Red Reagent and 0.15 U/ml xanthine oxidase (X2252-25UN, Sigma Aldrich) was added to the samples and incubated for 1 hour at room temperature. Finally, fluorescence of the samples was quantified in a plate reader with an excitation of 545 nm and absorption of 590 nm. For the continuous assay testing guanine and hypoxanthine directly with XO/horseradish peroxidase, 20 μl of each nucleobase was mixed with 170 μl XO/HRP reaction buffer in a microplate to final concentrations of 10, 100, and 500 μM. Fluorescence was measured as above at timepoints indicated in figure 3.

### Preparations of chemicals

XO (SIGMA, X2252-25UN) and HRP (SIGMA, P8375-5KU) were each resuspended in 25 mM HEPES (pH=7.4), 130 mM NaCl, 5 mM KCl, 50% glycerol, and 0.2 mg/ml BSA to a final concentration of 15 U/ml and 2.5 U/μl, respectively. Amplex Red reagent (Invitrogen, A12222) was resuspended in DMSO to a final concentration of 6 mM. Hypoxanthine (SIGMA, H9636) was dissolved in formic acid:H_2_O (v/v ratio of 2:1) to a concentration of 367 mM and was then diluted to 4.75, 0.95, and 0.095 mM in the PpnN reaction buffer. Guanine (SIGMA, G11950-10G) was resuspended in 15 mM KOH to a concentration of 10 mM and was then diluted to 4.75, 0.95, and 0.095 mM in the PpnN reaction buffer. GMP (G8377-500MG) was dissolved in 25 mM Tris-HCl (pH=8) to a concentration of 123 mM and was subsequently diluted into the PpnN reaction buffer to the desired concentrations. IMP (SIGMA, I4625-5G) was diluted into 127 mM HEPES to a final concentration of 50 mM. UMP (Alfa Aesar, A18601) and CMP (Alfa Aesar, J63376) were dissolved in MQ H_2_O to concentrations of 121 mM and 191 mM, respectively, and were then diluted into the PpnN reaction buffer to the desired concentrations. AMP (SIGMA, A2252-5G) was made by adding 0.191 g powder into 5 ml 25 mM Tris-HCl (pH=8), pH was adjusted by adding ca 50 μl 10 M NaOH and final volume was adjusted with MQ H_2_O to 10.5 ml.

## Acknowledgement

This study was supported by a Novo Nordisk Foundation Project Grant (NNF19OC0058331) to Y.E.Z.

---

**Table 1:** Bacterial strains used in this study.

| Strain | Relevant features | Reference |
| --- | --- | --- |
| YZ785 | Bl21 DE3 *ppnN::kan*, pCA24N-*ppnN* (*E. coli*), Kan, Cam | (2) |
| YZ693 | Bl21 DE3, pET24b-*ppnN*-6HIS (*S. typhimurium*), Kan | (5) |

## Notes

### Competing Interest Statement

The authors have declared no competing interest.

